# Atlas drug-eluting coronary stents inhibits neointimal hyperplasia in sheep modeling

**DOI:** 10.1101/2023.04.19.537461

**Authors:** Rasit Dinc, Halit Yerebakan

## Abstract

**Background:** Coronary artery disease (CAD) is one of the leading causes of mortality and morbidity worldwide. Many patients with CAD require mechanical revascularization. However, restenosis after minimally invasive intervention is a major problem for these patients. Fortunately, due to the controlled drug delivery properties of drug-eluting stents (DES), this problem seems to be significantly overcome. In this study, the pharmacodynamic and pharmacokinetic properties of Atlas sirolimus-eluting coronary stents coated with PLGA were evaluated.

**Materials and Methods:** This study was carried out in 20 non-atherosclerotic female sheep, divided into four groups that included 4 study and 1 control randomly assigned to each group. The animals in the study groups were stented with Atlas Drug-eluting coronary stents and the pharmacodynamic and pharmacokinetic properties were evaluated.

**Results:** A statistically important effectivity of sirolimus on the vascular endothelium was shown. With time, the decrease in sirolimus in blood samples was statistically significant. In the others, no clinically significant side effects were observed.

**Conclusions:** The results in this study showed a significant reduction in neointimal hyperplasia after experimental implantation of Atlas drug-eluting stents coated with PLGA polymer. Pharmacokinetic studies also showed that the stent did not release significant left sirolimus after 28 days.

## 1. Introduction

Coronary artery disease (CAD), the most common type of heart disease, continues to be the leading cause of mortality and morbidity worldwide (**1-3**).

Many patients with CAD require mechanical revascularization using coronary artery bypass grafting (CABG) or minimally invasive techniques (MIT) such as balloon and/or stent (**4**). Since the second half of the 20th century, minimally invasive techniques have revolutionized the field of interventional cardiology. Although more invasive CABG is superior in some groups of patients, they have become the preferred methods of revascularization (**4-6**). On the other hand, reocclusion of the coronary arteries due to thrombotic occlusion in acute stages or neointimal hyperplasia (NIH) in the longer periods following minimally invasive techniques has been a major problem after MIT, particularly after bare metal stents (BMS) (**7-9**). In the longer period re-occlusion process, multiple factors such as smooth muscle cell (SMC) migration, activation of some cytokines and growth factors, extracellular matrix formation, neutrophil/macrophage activation are involved. These events were depending on the injury caused by the stenting (**10**).

Drug-eluting stents (DES), which usually release sirolimus or paclitaxel, are now widely used to reduce the rate of restenosis (**9**). Compared to BMS, first-generation DES designed with nondegradable polymer coatings significantly reduced restenosis, but was associated with an increased risk of very late stent thrombosis. Later, second-generation DES designed with biocompatible more biocompatible durable polymers or bioabsorbable polymers have been developed using different platforms (**11**). Poly (lactic acid-coglycolic acid) (PLGA), a biodegradable and biocompatible polymer, is widely used in the coating of stents for the controlled delivery of various drugs to overcome these adverse effects (**5**).

The deployment of new devices requires testing for efficacy, feasibility, and safety (**4**). Sheep are one of the representative models for observing both the relationship between stent injury and neointimal growth, as well as the pharmacodynamic and pharmacokinetic properties of drugs (**12**). Several studies reported that drug-eluting polymer-coated stents significantly reduced stent thrombosis and had a good safety profile (**11, 13, 14**).

In this study, the pharmacodynamic and pharmacokinetic properties of Atlas sirolimus-eluting coronary stents coated with PLGA, which we recently produced as Invamed (Ankara, Turkey), were evaluated in sheep modeling before they were put into clinical use.

## 2. Materials and Methods

### 2.1. Ethical approval

The ethics committee approval was received from Kirikkale University Animal Experiments Local Ethics Committee (meeting date: 21.10.2020, meeting number: 2020/06, decision no: 40).

### 2.2. Ort of the study

The study was carried out at the Kirikkale University Scientific and Technological Research Application and Research Center (KUBTUAM).

### 2.3. Animals

This study was carried out on 20 female nonatherosclerotic Akkaraman ewes between 1 year old between 50-60 kg, provided through the Kirikkale University Huseyin Aytemiz Experimental Research and Application Laboratory. All animal experiments comply with the guidelines of the Directive 2010/63/EU of the European Parliament on the protection of animals used for scientific purposes and the study project.

The animals were divided into four groups. Each group included 4 study animals and 1 control animal randomly assigned to each group. The study was carried out for 26 weeks to determine the time course of the response of the vascular injury to stenting. For this reason, the animals were sacrificed and their stented arteries were harvested in the second postoperative week (Group 1), fourth week (Group 2), 12^th^ week (Group 3) and 26^th^ week (Group 4) (Table 1).

**Table 1.**
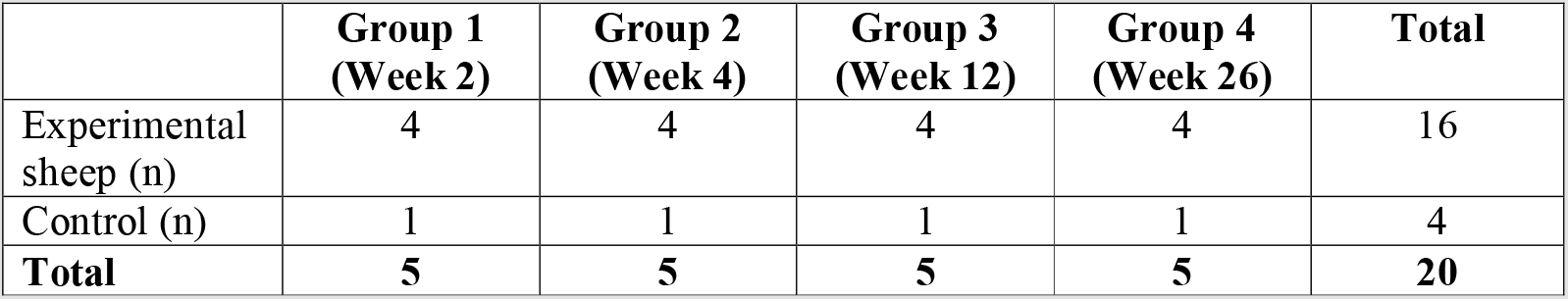
The numbers of sheep sacrificed (experimental and control) for the pharmacodynamic evaluation

### 2.4. Characteristics of the stent (Figure 1)

The stent brand name: Atlas Drug-Eluting Coronary Stent System (Invamed RD Global, Ankara, Turkey). The characteristics of the stent: Laser cut alloy with a wall thickness of 0.115 mm. Deployment: Balloon expandable. The stent coating: Deep-coating poly (lactic acid-coglycolic acid) (PLGA) containing sirolimus, 1µg/mm^2^, and auxiliary substances (Figure 1).

**Figure 1.**
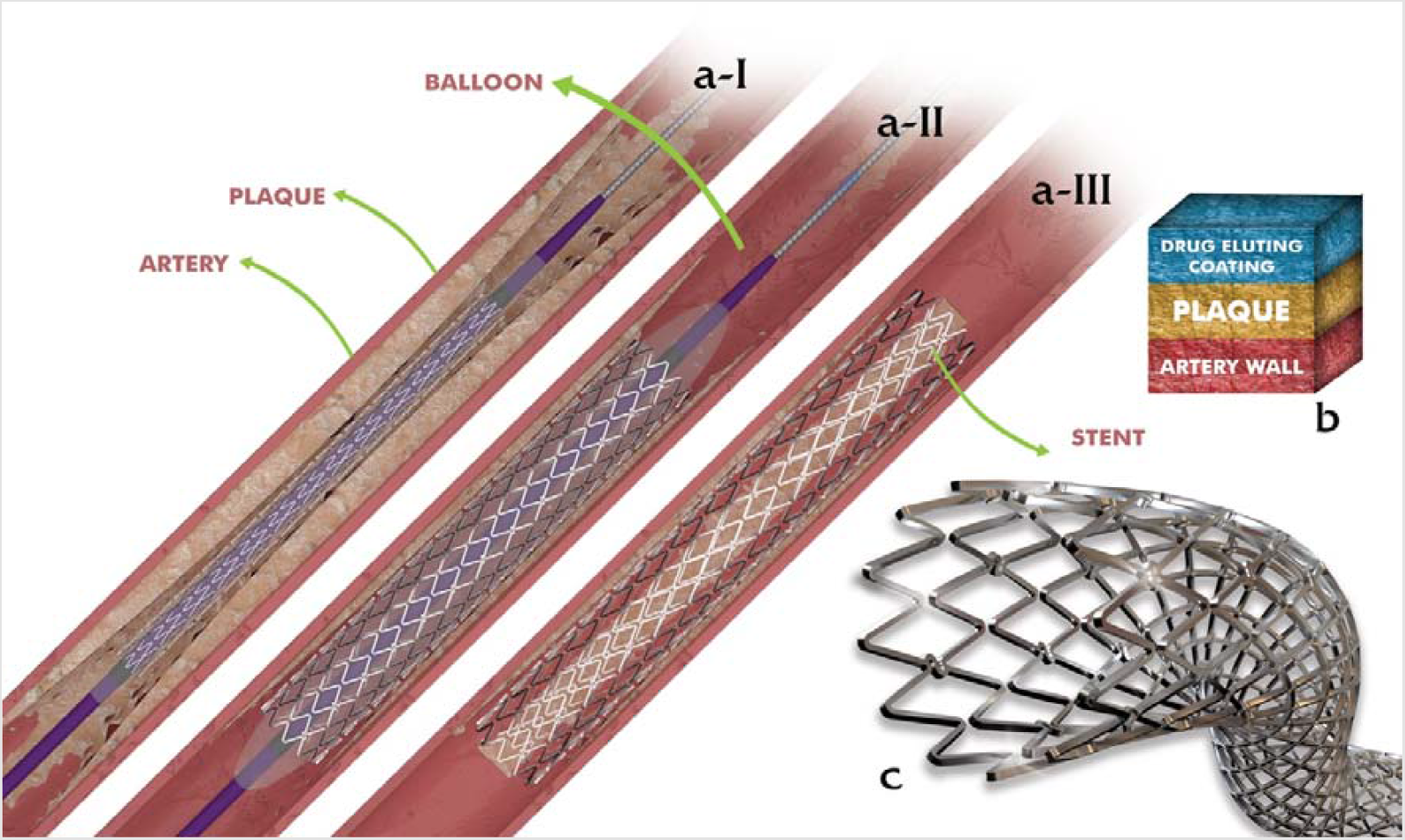
The structure of Atlas drug-eluting stent and its deployment in a diseased coronary artery. The illustrations in **a**) are **a-1**) just placed stent with a guidewire around the lesion site, **a-2**) balloon-expanded stent, and **a-3**) stent deployed after balloon removal. **b**. Cross-sectional view of the stent-inserted lesion site. **c**. Detailed structure of the stent

### 2.5. Stent placement into vessels

The animals were administered ticlopidine (antiplatelet) and sodium bemiparin (anticoagulant) 3 days before the intervention. All animals were injected with 0.25 mg / kg of diazepam and 2.5 mg / kg of ketamine for the intervention. An electrocardiogram was recorded with intravenous access. All stents were implanted under coronary angiography guidance using the femoral artery.

For stent implantation, the femoral artery in 1.1/1 extreme tension pressure was chosen in nonatherosclerotic arteries of sheep. The wire was sent to the left anterior descending artery and the right coronary artery, respectively, with a guide tube. A fixed stent pouch was sent along the guidewire. An expansion pressure of 10 atm for 30 seconds was applied with the balloon to place the stent in the proper place. Coronary angiography was performed before and after stent placement, as well as before the stent was sacrificed.

### 2.6. Pharmacokinetic evaluation

After implanting the stents, 0.5 ml of whole blood samples were collected from 16 sheep with a stent and one control in each planned time session (total 20 sessions): at 30^th^ minute; at 1^st^, 2^nd^, 4^th^, 8^th^, 12^th^, 24^th^ and 36^th^ hours; 2^nd^, 3^rd^, 4^th^ and 5 days; in 1st, 2^nd^, 3^rd^, 4^th^, 5^th^, 6^th^, 7^th^ and 8^th^ weeks. Blood samples were not collected from the animals in Group 1 after the fourth week and in Group 2 after the eighth week, since they were sacrificed in the second week and the fourth week, respectively.

The amount of sirolimus in the collected blood samples was detected by high performance liquid chromatography (HPLC) (Shimadzu, CTO-10AS) method according to the manufacturer’s instructions.

### 2.7. Pharmacodynamic Evaluation

For pharmacodynamic evaluation, integration of inflammation, tissue damage, and reendothelialization in the sacrificed animals in every planned time session for each group.

### 2.8. Histopathological analysis

After stent-implanted and normal sheep were sacrificed, coronary artery tissue biopsy specimens were stained with standard hematoxylin-eosin (HE, Sigma-Aldrich, USA) after fixation in 10% buffered formalin according to previously described (**15**). All stained samples were examined under a microscope (Olympus BX51; Olympus C&S, Japan) and their microphotographs were taken (DP25; Olympus C&S, Japan).

The proximal, medial and distal parts of the intimal layer of the stented area of the artery, which includes the internal elastic lamina and endothelium, were evaluated using a 10X objective of the microscope. Histopathological findings were evaluated according to endothelialization criteria, presence of thrombosis, and endothelial and intimal thickening. The data obtained were scored according to the reports described previously (**16**) (Table 2).

**Table 2.**
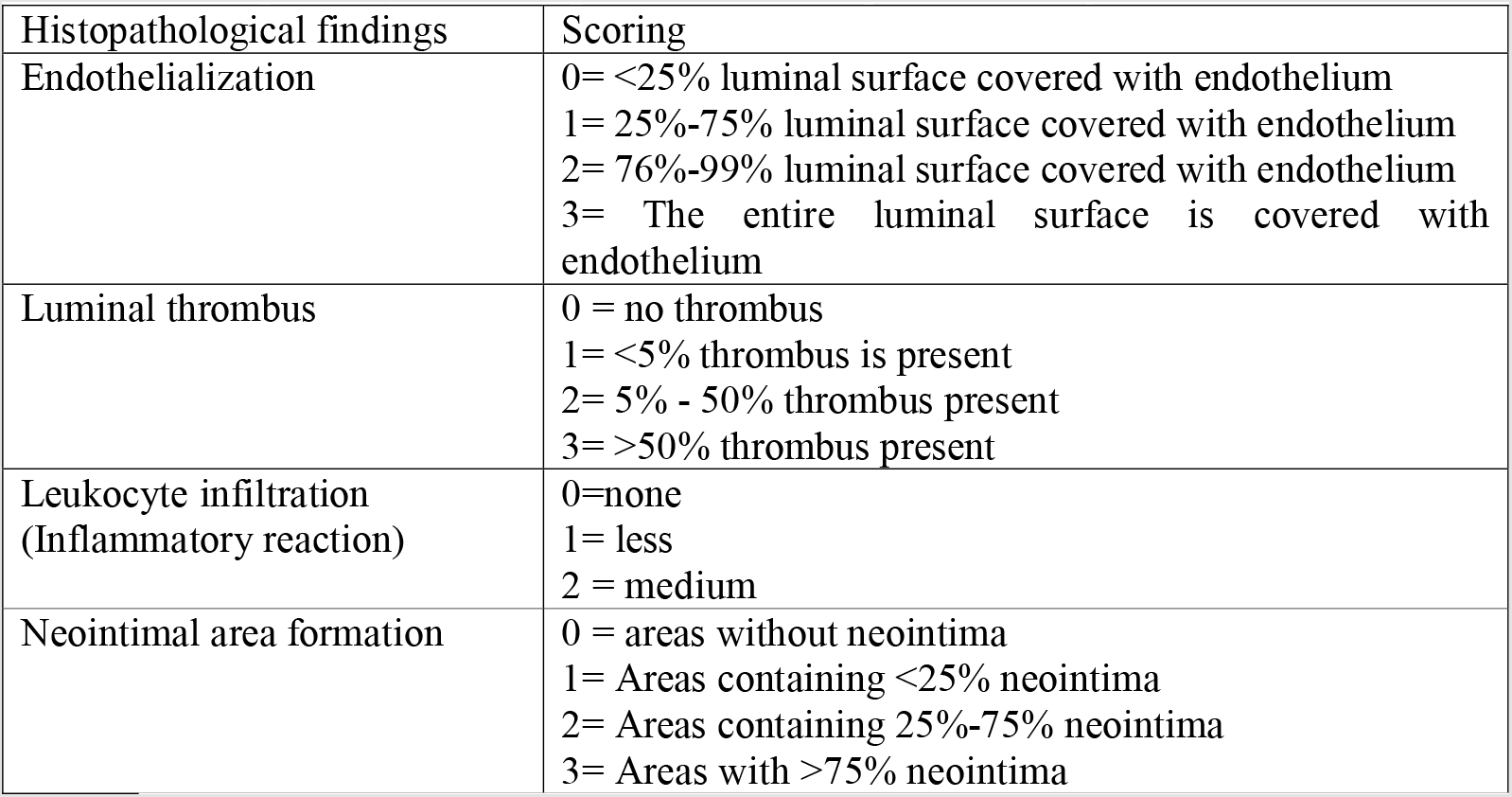
Histopathological scoring of coronary artery tissue biopsy specimens

| Histopathological findings | Scoring |
| --- | --- |
| Endothelialization | 0= <25% luminal surface covered with endothelium<br>1= 25%-75% luminal surface covered with endothelium<br>2= 76%-99% luminal surface covered with endothelium<br>3= The entire luminal surface is covered with endothelium |
| Luminal thrombus | 0 = no thrombus<br>1= <5% thrombus is present<br>2= 5% - 50% thrombus present<br>3= >50% thrombus present |
| Leukocyte infiltration<br>(Inflammatory reaction) | 0=none<br>1= less<br>2 = medium |
| Neointimal area formation | 0 = areas without neointima<br>1= Areas containing <25% neointima<br>2= Areas containing 25%-75% neointima<br>3= Areas with >75% neointima |

The proximal, medial and distal parts of the stented area were stained with Verhoeff-van Gieson (VVG) dyes and the coronal sections were measured using a 10X objective. Quantitative histomorphometric measurements were completed by taking the averages with the lmageJ program (v1.46r; USA). The difference between the inner elastic lamina area and the cross-sectional lumen area was determined as the neointimal area. The percentage of vascular lumen narrowing is calculated using the formula “[1-luminal area/IEL area]X100”.

### 2.9. Statistical analysis

For data analysis, IBM SPSS Version 21 and the MedCalc statistical package program (MedCalc Software, Ostend, Belgium) were used. Parametric tests were used without the normality test due to the compatibility of the central limit theorem (**17**). The mean and standard deviation values were used for the statistics of continuous data in the scales.

The repeated ANOVA test statistic was used to compare the mean of more than two independent measurements and the one-way ANOVA test statistic was used to compare the mean of continuous measurements in more than two independent groups. In case of difference, it was evaluated with Bonferroni and Tukey statistics as a post hoc test.

The statistical significance level of the data was taken as p<0.05.

## 3. Results

An animal in groups 3 (no. 12) and an animal in 4 (no. 14) died after implantation. Therefore, these two animals were excluded from the evaluation. In others, no significant adverse events were clinically observed during the entire follow-up period. 14 implanted animals were sacrificed at different predetermined time intervals. Thrombus formation was observed in 2 of the vessel samples, 1 from group-3 and 1 from group-4, in the histopathological evaluation.

### 3.1. Macroscopic Results

Stent-induced tissue integrity deterioration and stenosis were macroscopically not observed in any of the stented vessels (Figure 2).

**Figure 2:**
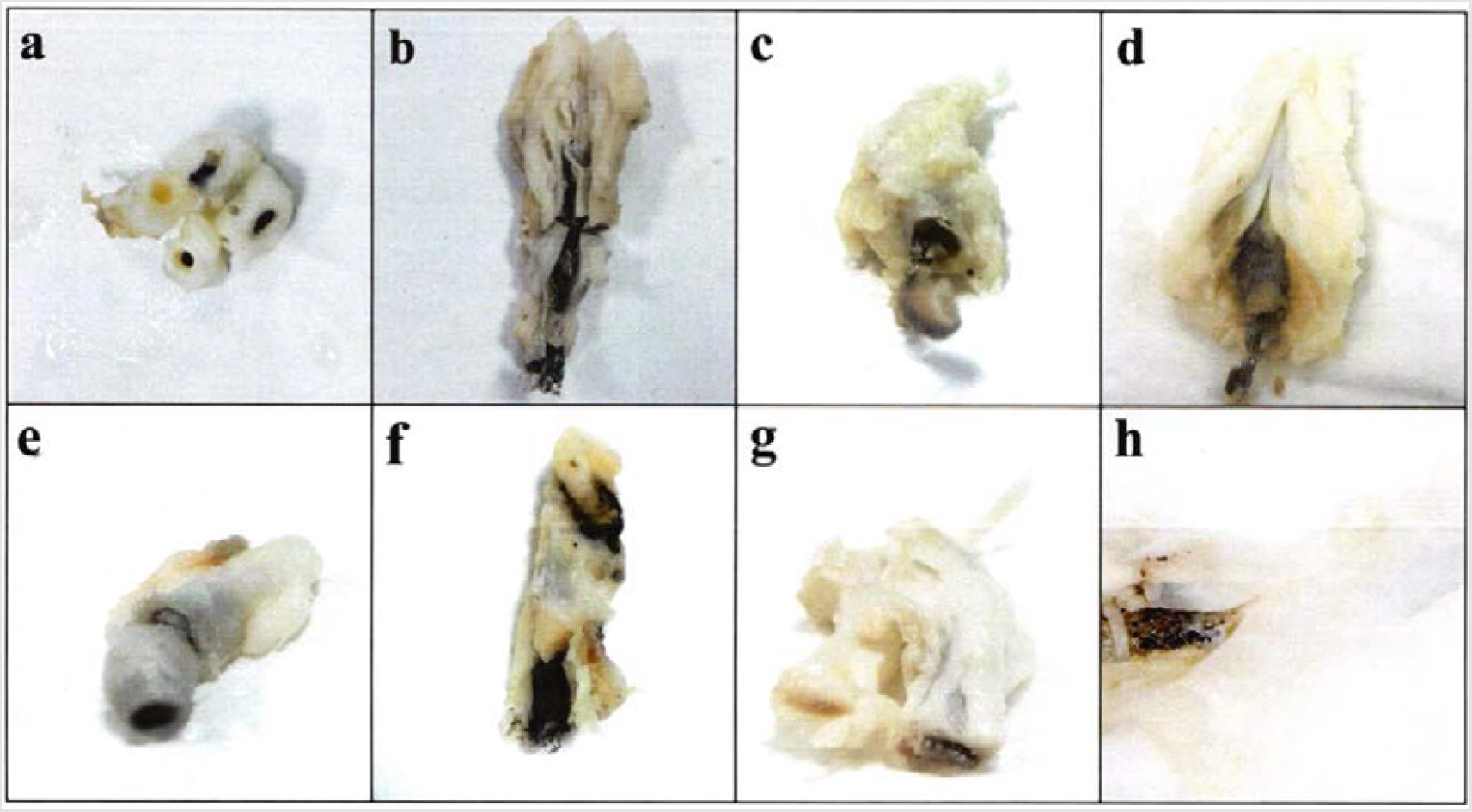
Macroscopic pictures of the stented arteries. a-b. Group-1 sagittal and longitudinal sections, respectively. c-d. Group-2 sagittal and longitudinal section, respectively. e-f. Group-3 sagittal and longitudinal sections, respectively. g-h. Group 4 sagittal and longitudinal section, respectively.

### 3.2. Histopathological Results

Compared to the control group, reendothelialization was observed at a rate of approximately 90% in vessel samples from Group 1, Group 2, and Group 4 and the surface was covered with endothelium (Table 3, Figure 3 B-C-D-E-H-I). In Group 2, except for sample 7, it was observed that the reendothelialization in the vessel was not fully formed and the reendothelialization rate remained in the range of 70% to 90% (Table 3, Figure 3 F and G). Recanalized microthrombus (score 3= >50%) was found in 1 sample of vessels in Group 3 (Figure 3 F), while mild accumulation of fibrinogen and erythrocytes was found in 1 sample of vessels in Group 4 (Figure 3 H).

**Table 3.**
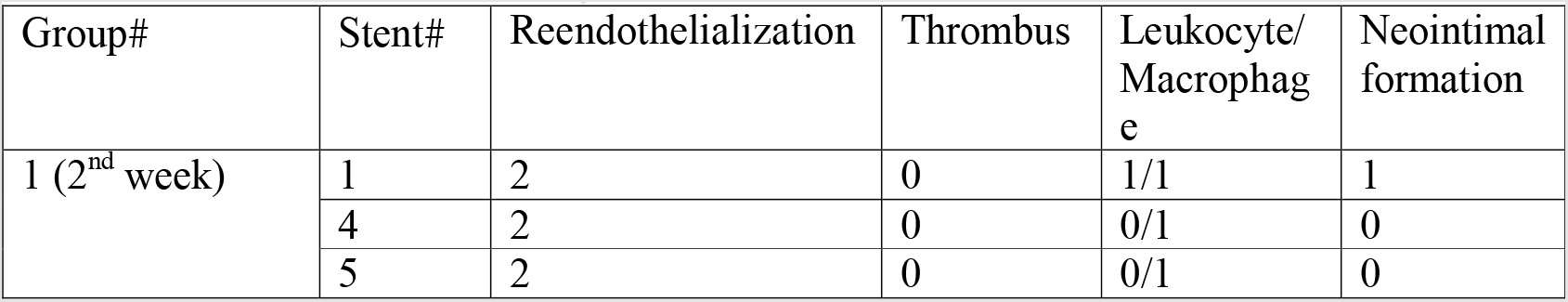
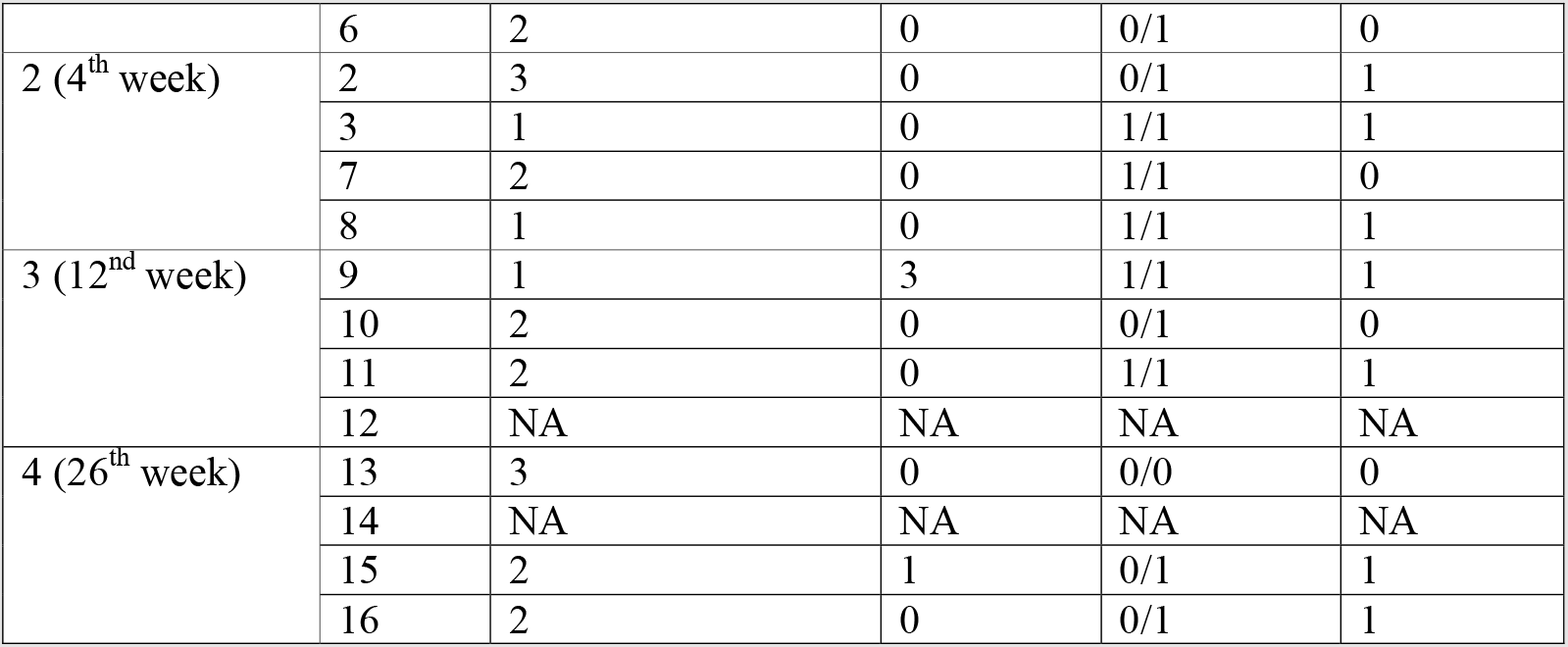
Histopathological scoring of the stented arteries

**Figure 3:**
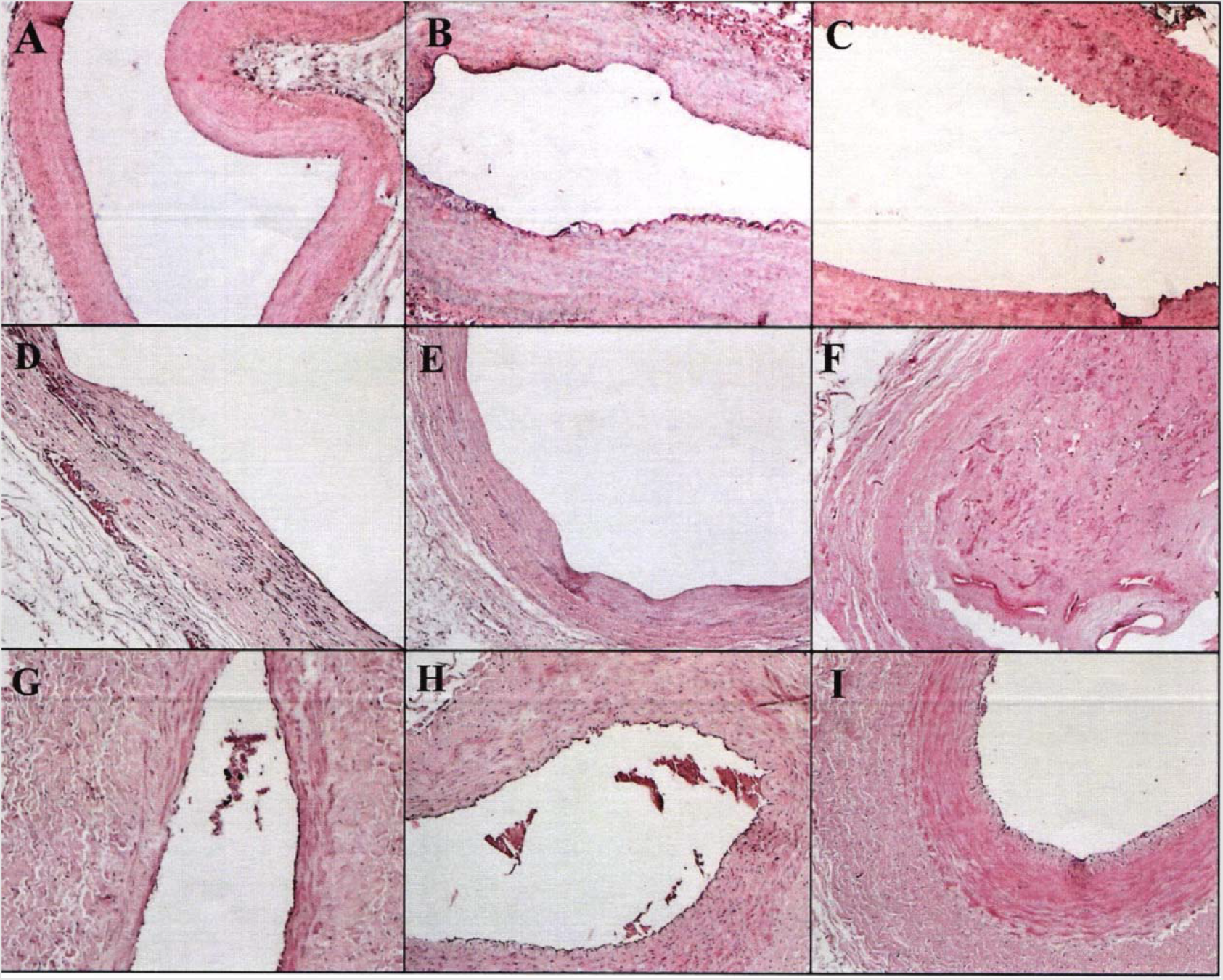
HE-stained microscopic pictures of vessel samples from control, Groups 1,2,3 and 4. **A (Control group**). Complete endothelialization (score 3=100%), no thrombus, and no inflammatory infiltration; 4x magnification. **B-C (Group-1, Week-2**). Endothelial formation is complete approximately 90% (score 2 = 76-99%), mild intimal thickening with mild macrophage infiltration; 4x magnification. **E-D (Group-2, Week-4**). Endothelial formation is largely complete, mild intimal/medial thickening with mild to moderate macrophage infiltration; 4x and 10x, respectively. **F-G (Group-3, Week-12**). Endothelial formation is complete 70-90% (score 1= 50-75%) with mild intimal/medial thickening with mild to moderate macrophage infiltration; 4x and 10x, respectively. The presence of more than 50% (score 3= >50%) recanalized thrombus in F and the presence of erythrocyte/inflammatory cells in G are notable. **H-I (Group-4, Week 26**). Endothelial formation is largely complete, intimal / medial thickening with mild macrophage and leukocyte infiltration. Inflammatory cell and erythrocyte infiltration in the lumen of the vessel and intimal surface is remarkable; 10x and 4x, respectively.

Mild neutrophil leukocyte and macrophage infiltration in vessel samples in Groups 1 and 4 and, consequently, mild thickening in the intimal and medial regions. In Groups 2 and 3, macrophage infiltration of varying severity from mild to moderate and again mild leukocyte infiltration were found. Thus, both medial and intimal thickenings were recorded in Groups 2 and 3 to be higher compared to the control group and Groups 1 and 4.

A neointimal area of <10% was observed in 8 of the stented vessels. No proximal or distal end stenosis was found in any of the vessels.

### 3.2. Histomorphometric evaluation

There were statistically significant differences between the means of the measurements of the IEL, EEL, the medial area, the neointimal area, and the luminal area according to the groups (p†<0.05) (Table 4).

**Table 4.** Histomorphometric scoring of the stented arteries and comparison according to groups.

| Group# | Stent# | IEL (mm<sup>2</sup>) | EEL (mm<sup>2</sup>) | Medial area (mm<sup>2</sup>) | Neointimal area (mm<sup>2</sup>) | Luminal Area (mm<sup>2</sup>) |
| --- | --- | --- | --- | --- | --- | --- |
| 1 (2<sup>nd</sup> week) | 1 | 5,90 ± 0,85 | 7,61 ± 0,66 | 1,71 ± 0,51 | 3,21 ± 0,61 | 2,69 ± 1,07 |
|  | 4 | 5,88 ± 1,09 | 7,59 ± 0,81 | 1,71 ± 0,27 | 3,25 ± 0,25 | 2,63 ± 0,67 |
|  | 5 | 5,89 ± 0,77 | 7,56 ± 0,95 | 1,67 ± 0,09 | 3,21 ± 0,39 | 2,68 ± 1,03 |
|  | 6 | 5,91 ± 0,92 | 7,63 ± 0,49 | 1,72 ± 0,14 | 3,27 ± 0,87 | 2,64 ± 0,84 |
|  | **Mean±Std.** | **5,89±0,01** | **7,59±0,03** | **1,71±0,02** | **3,23±0,03** | **2,66±0,03** |
| 2 (4<sup>th</sup> week) | 2 | 4,84 ± 0,55 | 7,89 ± 1,06 | 3,05 ± 0,23 | 2,53 ± 1,06 | 2,31 ± 0,93 |
|  | 3 | 4,96 ± 0,38 | 7,93 ± 0,35 | 2,43 ± 0,17 | 2,68 ± 0,88 | 2,28 ± 0,87 |
|  | 7 | 4,93 ± 0,69 | 7,91 ± 0,94 | 2,98 ± 0,36 | 2,66 ± 0,64 | 2,27 ± 0,36 |
|  | 8 | 4,88 ± 0,95 | 7,96 ± 1,01 | 3,08 ± 0,98 | 2,58 ± 0,96 | 2,30 ± 0,71 |
|  | **Mean±Std.** | **4,91±0,05** | **7,92±0,03** | **2,89±0,31** | **2,62±0,06** | **2,29±0,02** |
| 3 (12<sup>nd</sup> week) | 9 | 3,76 ± 1,45 | 8,72 ± 2,30 | 4,96 ± 0,21 | 2,49 ± 0,98 | 1,27 ± 0,87 |
|  | 10 | 3,75 ± 1,08 | 8,55 ± 1,58 | 4,80 ± 0,62 | 2,51 ± 0,65 | 1,24 ± 0,66 |
|  | 11 | 3,69 ± 1,35 | 8,63 ± 1,67 | 4,94 ± 0,74 | 2,39 ± 0,33 | 1,30 ± 0,82 |
|  | 12 | NA | NA | NA | NA | NA |
|  | **Mean±Std.** | **3,73±0,04** | **8,63±0,08** | **4,9±0,09** | **2,46±0,06** | **1,27±0,03** |
| 4 (26<sup>th</sup> week) | 13 | 5,98 ± 0,41 | 7,99 ± 1,45 | 2,01 ± 0,29 | 4,24 ± 0,65 | 1,74 ± 0,02 |
|  | 14 | NA | NA | NA | NA | NA |
|  | 15 | 5,84 ± 0,59 | 8,01 ± 1,56 | 2,17 ± 0,32 | 4,12 ± 0,74 | 1,72 ± 0,11 |
|  | 16 | 5,89 ± 0,27 | 7,96 ± 1,06 | 2,07 ± 0,58 | 4,14 ± 0,08 | 1,75 ± 0,28 |
|  | **Mean±Std.** | **5,91±0,07** | **7,98±0,2** | **2,08±0,08** | **4,17±0,06** | **1,74±0,01** |
| p value* |  | <0,001 | <0,001 | <0,001 | <0,001 | <0,001 |

\* The p-value was calculated according to the mean of the subjects in the groups using the one-way ANOVA test.
**Abbreviations** IEL: Internal endothelial lumen, EEL: External endothelial lumen.

### 3.3. Results of the Pharmacodynamic Evaluation

There were statistically significant differences between the mean diameter of the vessel before stent implantation, MLD after stent implantation, RVD, MLD before sacrifice, luminal loss and restenosis ratio according to the groups (p†<0.05) (Table 5).

**Table 5.** Effect of the stent on the arterial lumen and comparison according to groups.

| Group# | Stent# | Vessel diameter prior to stent | MLD after stent | RVD | MLD before sacrificing | Luminal | Restenosis |
| --- | --- | --- | --- | --- | --- | --- | --- |

|  |  | implantation<br>(mm) | implantation<br>(mm) | (mm) | (mm) | loss<br>(mm)* | ratio<br>(%) |
| --- | --- | --- | --- | --- | --- | --- | --- |
| 1 (2<sup>nd</sup> week) | 1 | 2,54 ± 0,15 | 2,97 ± 0,18 | 3,10 ± 0,18 | 1,52 ± 0,17 | 1,45 ± 0,21 | 50,96 ± 3,51 |
|  | 4 | 2,61 ± 0,12 | 3,11 ± 0,15 | 3,16 ± 0,11 | 1,50 ± 0,21 | 1,61 ± 0,27 | 52,53 ± 3,71 |
|  | 5 | 2,59 ± 0,18 | 2,92 ± 0,16 | 2,59 ± 0,16 | 1,50 ± 0,23 | 1,42 ± 0,26 | 42,08 ± 6,63 |
|  | 6 | 2,58 ± 0,16 | 3,04 ± 0,12 | 2,63 ± 0,15 | 1,54 ± 0,15 | 1,50 ± 0,24 | 41,44 ± 5,76 |
| **Mean±Std.** |  | **2,58±0,02** | **3,01±0,08** | **2,87±0,3** | **1,51±0,02** | **1,49±0,08** | **46,75±5,81** |
| 2 (4<sup>th</sup> week) | 2 | 2,51 ± 0,13 | 2,68 ± 0,24 | 2,90 ± 0,15 | 1,76 ± 0,35 | 0,92 ± 0,41 | 41,54 ± 8,44 |
|  | 3 | 2,56 ± 0,19 | 2,99 ± 0,27 | 3,05 ± 0,19 | 1,81 ± 0,11 | 1,18 ± 0,43 | 40,65 ± 10,32 |
|  | 7 | 2,54 ± 0,10 | 2,87 ± 0,32 | 3,02 ± 0,34 | 1,78 ± 0,17 | 1,09 ± 0,28 | 41,06 ± 6,38 |
|  | 8 | 2,57 ± 0,14 | 2,95 ± 0,21 | 3,07 ± 0,13 | 1,80 ± 0,22 | 1,15 ± 0,69 | 41,36 ± 9,41 |
| **Mean±Std.** |  | **2,55±0,03** | **2,87±0,14** | **3,01±0,08** | **1,79±0,02** | **1,09±0,11** | **41,15±0,39** |
| 3 (12<sup>nd</sup> week) | 9 | 2,59 ± 0,18 | 2,92 ± 0,16 | 2,61 ± 0,16 | 1,50 ± 0,18 | 1,42 ± 0,26 | 45,10 ± 6,47 |
|  | 10 | 2,62 ± 0,08 | 2,95 ± 0,12 | 2,64 ± 0,19 | 1,56 ± 0,24 | 1,39 ± 0,45 | 40,91 ± 7,26 |
|  | 11 | 2,64 ± 0,23 | 2,99 ± 0,10 | 2,67 ± 0,09 | 1,61 ± 0,18 | 1,38 ± 0,22 | 39,70 ± 4,05 |
|  | 12 | NA | NA | NA | NA | NA | NA |
| **Mean±Std.** |  | **2,62±0,02** | **2,95±0,03** | **2,64±0,03** | **1,56±0,06** | **1,39±0,02** | **41,91±2,83** |
| 4 (26<sup>th</sup> week) | 13 | 2,79 ± 0,13 | 3,31 ± 0,14 | 1,40 ± 0,06 | 2,83 ± 0,18 | 2,86 ± 0,10 | 1,49 ± 0,45 |
|  | 14 | NA | NA | NA | NA | NA | NA |
|  | 15 | 2,69 ± 0,17 | 3,28 ± 0,18 | 1,42 ± 0,02 | 2,85 ± 0,16 | 2,87 ± 0,07 | 1,50 ± 0,41 |
|  | 16 | 2,73 ± 0,09 | 3,30 ± 0,21 | 1,39 ± 0,05 | 2,86 ± 0,25 | 2,89 ± 0,13 | 1,53 ± 0,34 |
| **Mean±Std.** |  | **2,73±0,05** | **3,29±0,01** | **1,41±0,01** | **2,85±0,01** | **2,87±0,02** | **1,51±0,02** |
| p_value* |  | <0,001 | <0,001 | <0,001 | <0,001 | <0,001 | <0,001 |

\* The p-value was calculated according to the mean of the subjects in the groups using the one-way ANOVA test.
**Abbreviations** MLD: Minimal lumen diameter, RVD: Reference vessel diameter.

### 3.4. Results of the Pharmacokinetic Evaluation

As shown in Tables 6 and Figure 4, the concentration of sirolimus in the blood peaked within the first 2 hours after stent implantation. Its levels gradually decreased from 4 hours to 8 weeks.

**Table 6.**
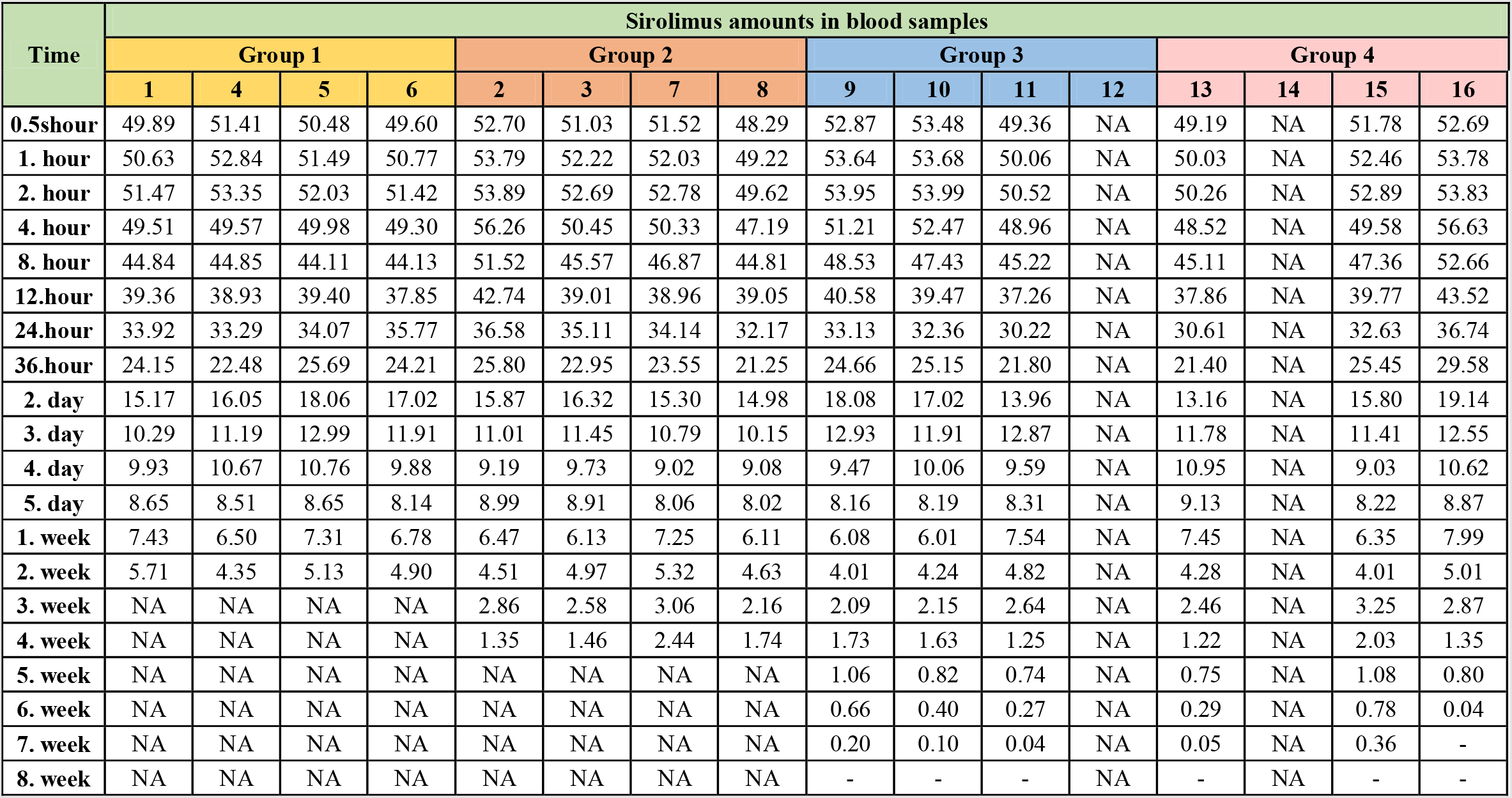
Time course of sirolimus concentration (ng/mL)

| Time | Sirolimus amounts in blood samples |  |  |  |  |  |  |  |  |  |  |  |  |  |  |  |
| --- | --- | --- | --- | --- | --- | --- | --- | --- | --- | --- | --- | --- | --- | --- | --- | --- |
|  | Group 1 |  |  |  | Group 2 |  |  |  | Group 3 |  |  |  | Group 4 |  |  |  |
|  | 1 | 4 | 5 | 6 | 2 | 3 | 7 | 8 | 9 | 10 | 11 | 12 | 13 | 14 | 15 | 16 |
| **0.5hour** | 49.89 | 51.41 | 50.48 | 49.60 | 52.70 | 51.03 | 51.52 | 48.29 | 52.87 | 53.48 | 49.36 | NA | 49.19 | NA | 51.78 | 52.69 |
| **1. hour** | 50.63 | 52.84 | 51.49 | 50.77 | 53.79 | 52.22 | 52.03 | 49.22 | 53.64 | 53.68 | 50.06 | NA | 50.03 | NA | 52.46 | 53.78 |
| **2. hour** | 51.47 | 53.35 | 52.03 | 51.42 | 53.89 | 52.69 | 52.78 | 49.62 | 53.95 | 53.99 | 50.52 | NA | 50.26 | NA | 52.89 | 53.83 |
| **4. hour** | 49.51 | 49.57 | 49.98 | 49.30 | 56.26 | 50.45 | 50.33 | 47.19 | 51.21 | 52.47 | 48.96 | NA | 48.52 | NA | 49.58 | 56.63 |
| **8. hour** | 44.84 | 44.85 | 44.11 | 44.13 | 51.52 | 45.57 | 46.87 | 44.81 | 48.53 | 47.43 | 45.22 | NA | 45.11 | NA | 47.36 | 52.66 |
| **12.hour** | 39.36 | 38.93 | 39.40 | 37.85 | 42.74 | 39.01 | 38.96 | 39.05 | 40.58 | 39.47 | 37.26 | NA | 37.86 | NA | 39.77 | 43.52 |
| **24.hour** | 33.92 | 33.29 | 34.07 | 35.77 | 36.58 | 35.11 | 34.14 | 32.17 | 33.13 | 32.36 | 30.22 | NA | 30.61 | NA | 32.63 | 36.74 |
| **36.hour** | 24.15 | 22.48 | 25.69 | 24.21 | 25.80 | 22.95 | 23.55 | 21.25 | 24.66 | 25.15 | 21.80 | NA | 21.40 | NA | 25.45 | 29.58 |
| **2. day** | 15.17 | 16.05 | 18.06 | 17.02 | 15.87 | 16.32 | 15.30 | 14.98 | 18.08 | 17.02 | 13.96 | NA | 13.16 | NA | 15.80 | 19.14 |
| **3. day** | 10.29 | 11.19 | 12.99 | 11.91 | 11.01 | 11.45 | 10.79 | 10.15 | 12.93 | 11.91 | 12.87 | NA | 11.78 | NA | 11.41 | 12.55 |
| **4. day** | 9.93 | 10.67 | 10.76 | 9.88 | 9.19 | 9.73 | 9.02 | 9.08 | 9.47 | 10.06 | 9.59 | NA | 10.95 | NA | 9.03 | 10.62 |
| **5. day** | 8.65 | 8.51 | 8.65 | 8.14 | 8.99 | 8.91 | 8.06 | 8.02 | 8.16 | 8.19 | 8.31 | NA | 9.13 | NA | 8.22 | 8.87 |
| **1. week** | 7.43 | 6.50 | 7.31 | 6.78 | 6.47 | 6.13 | 7.25 | 6.11 | 6.08 | 6.01 | 7.54 | NA | 7.45 | NA | 6.35 | 7.99 |
| **2. week** | 5.71 | 4.35 | 5.13 | 4.90 | 4.51 | 4.97 | 5.32 | 4.63 | 4.01 | 4.24 | 4.82 | NA | 4.28 | NA | 4.01 | 5.01 |
| **3. week** | NA | NA | NA | NA | 2.86 | 2.58 | 3.06 | 2.16 | 2.09 | 2.15 | 2.64 | NA | 2.46 | NA | 3.25 | 2.87 |
| **4. week** | NA | NA | NA | NA | 1.35 | 1.46 | 2.44 | 1.74 | 1.73 | 1.63 | 1.25 | NA | 1.22 | NA | 2.03 | 1.35 |
| **5. week** | NA | NA | NA | NA | NA | NA | NA | NA | 1.06 | 0.82 | 0.74 | NA | 0.75 | NA | 1.08 | 0.80 |
| **6. week** | NA | NA | NA | NA | NA | NA | NA | NA | 0.66 | 0.40 | 0.27 | NA | 0.29 | NA | 0.78 | 0.04 |
| **7. week** | NA | NA | NA | NA | NA | NA | NA | NA | 0.20 | 0.10 | 0.04 | NA | 0.05 | NA | 0.36 | - |
| **8. week** | NA | NA | NA | NA | NA | NA | NA | NA | - | - | - | NA | - | NA | - | - |

**Figure 4.**
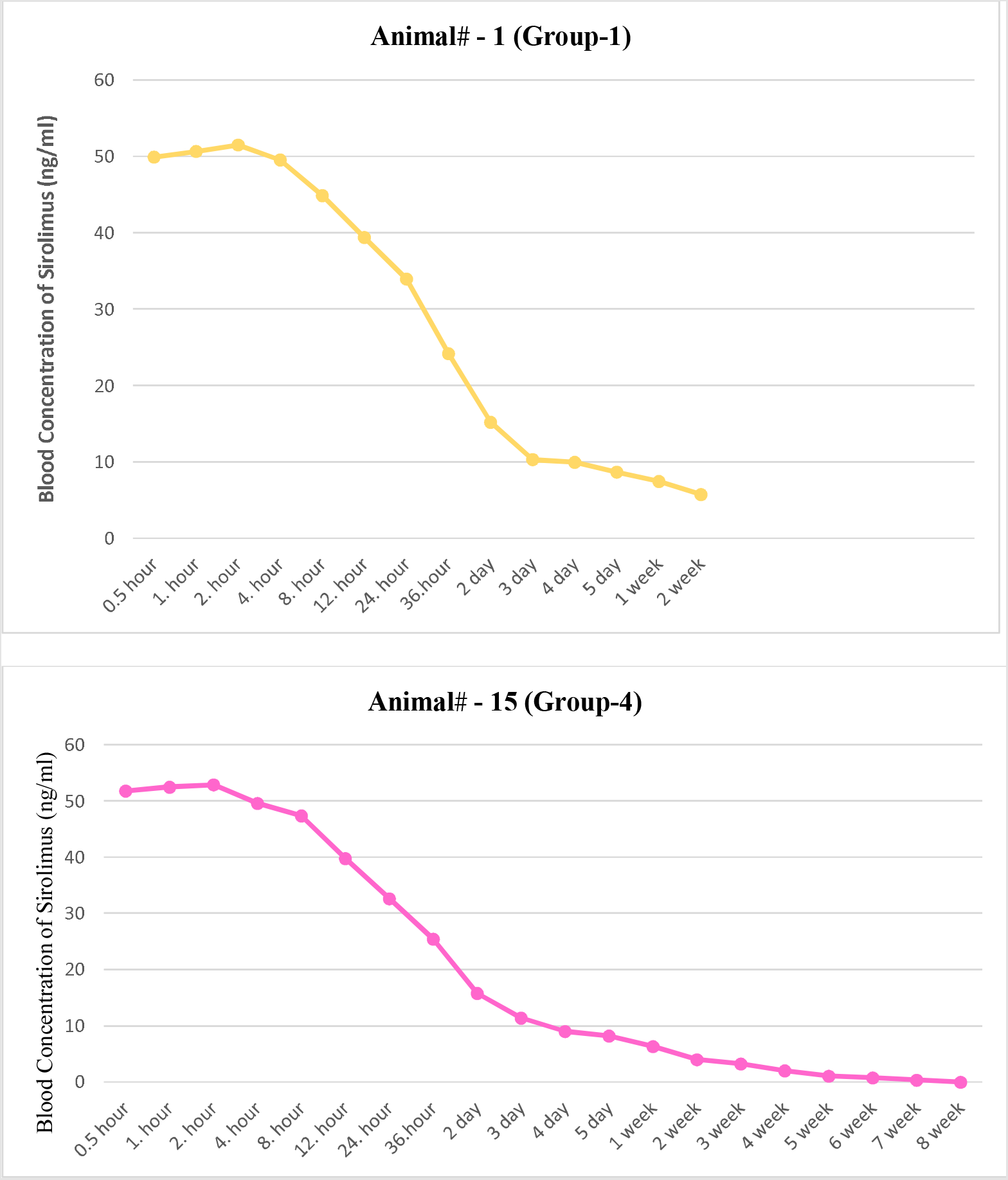
Time course plot of sirolimus concentration (ng/mL). *Representative samples are from Groups 1 and 4.

As we can see in Table 7 and Figure 5, there was statistically no difference in the mean sirolimus concentration between the groups at each time (p†>0.05). While the change in sirolimus concentration over time was significant in all groups (p††<0.005).

**Table 7.**
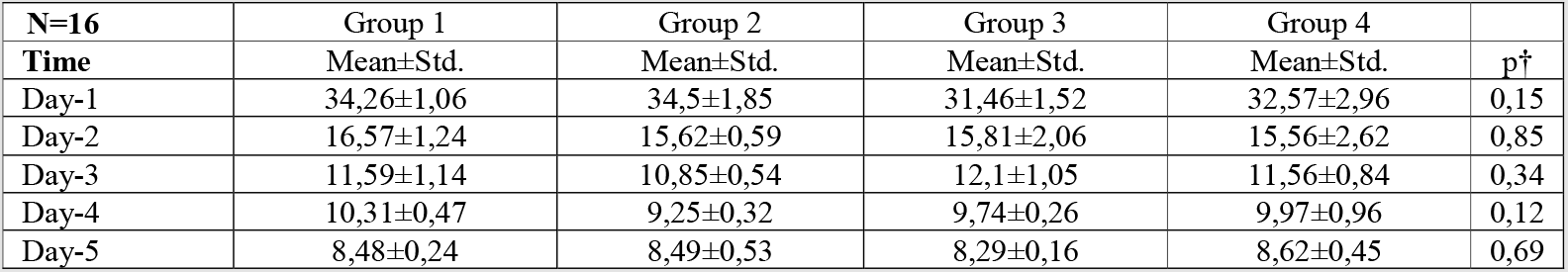
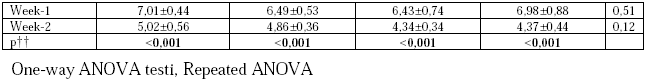
Comparison of sirolimus concentration by groups and time.

| N=16 | Group 1 | Group 2 | Group 3 | Group 4 |  |
| --- | --- | --- | --- | --- | --- |
| Time | Mean±Std. | Mean±Std. | Mean±Std. | Mean±Std. | $p^{\dagger}$ |
| Day-1 | 34,26±1,06 | 34,5±1,85 | 31,46±1,52 | 32,57±2,96 | 0,15 |
| Day-2 | 16,57±1,24 | 15,62±0,59 | 15,81±2,06 | 15,56±2,62 | 0,85 |
| Day-3 | 11,59±1,14 | 10,85±0,54 | 12,1±1,05 | 11,56±0,84 | 0,34 |
| Day-4 | 10,31±0,47 | 9,25±0,32 | 9,74±0,26 | 9,97±0,96 | 0,12 |
| Day-5 | 8,48±0,24 | 8,49±0,53 | 8,29±0,16 | 8,62±0,45 | 0,69 |
| Week-1 | 7,01±0,44 | 6,49±0,53 | 6,43±0,74 | 6,98±0,88 | 0,51 |
| Week-2 | 5,02±0,56 | 4,86±0,36 | 4,34±0,34 | 4,37±0,44 | 0,12 |
| p†† | <0,001 | <0,001 | <0,001 | <0,001 |  |

One-way ANOVA testi, Repeated ANOVA

**Figure 5.**
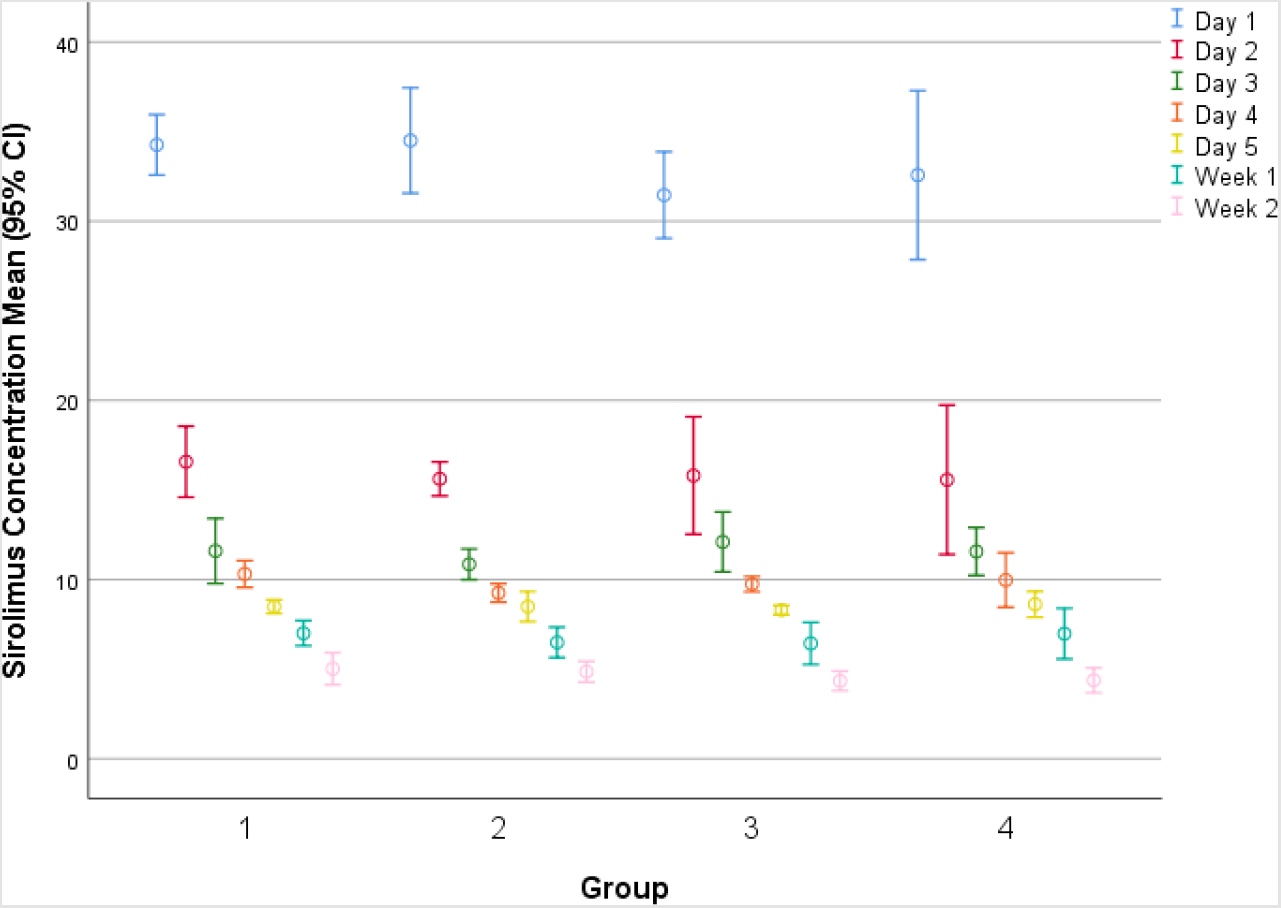
Comparison of sirolimus concentration by groups and time.

## Discussions

Stents have foreign body potential for vascular endothelium and circulating blood cells. The deployment of stents triggers thrombus formation, endothelial hyperreactivity, inflammatory events, and smooth muscle cell migration/proliferation. These processes eventually result in neointimal hyperplasia that contributes to restenosis after stent deployment (**15**). Permanent polymers in drug-eluting stents used to overcome restenosis adversely affect the stented vascular restoration and prolong the inflammatory reaction process. Eventually, they contribute to late and very late thrombosis (**9, 18**). When a biodegradable polymer is used, the drug is absorbed from the stent surface after elution, leaving only the neointima- and endothelium-coated metal stent without further irritating the arterial wall. This improves arterial vascular restoration and reduces the inflammatory reaction. It also reduces the need for long-term antiplatelet therapy (**18**).

Our study was conducted in sheep modeling. Because sheep are considered a suitable modeling for cardiovascular intervention due to their ease of handling, size, and vascular anatomy similar to those of humans (**19**). Sirolimus, on the other hand, is one of the widely used drugs to prevent restenosis (**9**). In target cells, it binds to the FK506 binding protein-12 and ultimately regulates downstream signaling pathways involved in cell survival (**20**). Consequently, the drug prevents restenosis by inhibiting cell cycles, including smooth muscle cells and inflammatory cells, both of which play an important role in stenosis (**10**,**21**). Some studies also confirm pathologically that these drugs delay arterial healing compared to the control group (**22**). Additionally, the multilayer absorbable PLGA coating completely degrades in vitro in approximately 8 weeks, immediately after complete elution of the drug (**4**,**23**). In addition to these, there are many studies demonstrating that biodegradable polymer coated sirolimus-eluting stents are both safe and feasible in preventing restenosis in human or experimental animal models (**24-27**). The metal structure of Atlas drug-eluting stents enables the stent to be prepared with a very thin support structure and allows radioopacity (**8**,**28**). Its biodegradable and biocompatible PLGA polymer has been extensively studied for the controlled delivery of several drugs and its characteristics are well known (**5**).

In the pharmacodynamic evaluation, which shows the effectivity of sirolimus on the endothelium, there were statistical differences for all parameters evaluated between the groups (p<0.001). Medial and intimal thickening was observed more in Groups 2 and 3 compared to Groups 1 and 4. This has been evaluated as due to the fact that the effect of sirolimus has not reached the highest level in Group 1, and that the sirolimus level has decreased too much in Group 4.

We did not follow the result after 26 weeks. However, it can be predicted that reendothelialization will develop on the stent strut because of the decrease in irritation of the vessel wall as a result of the almost disappearance of the effects of the polymer and drug. Because our pharmacokinetic studies have shown that the concentration of sirolimus in the blood cannot be determined after the eighth week after the implantation of the stents.

Over time, the decrease in sirolimus in blood samples was statistically significant in all groups (p<0.001). On the other hand, there were no statistical differences in the mean sirolimus concentration between the groups at each measurement time (p>0.05). These results show that Atlas drug elimination provides a consistent outcome in terms of drug release.

The general safety and efficacy results of this study were evaluated in accordance with previous reports. Two of the 14 animals that survived developed thrombus formation. No clinically significant side effects were observed.

## Study limitations

There were some limitations to the study. The sheep animal model may not fully reflect human characteristics to assess restenosis after experimental coronary interventions. Due to the short duration of the experiment, it is not possible to observe the consequences of long-term effects in animal models.

## Conclusions

The results in this study showed a significant reduction in neointimal hyperplasia after experimental implantation of Atlas drug-eluting stents coated with PLGA polymer. Pharmacokinetic studies also showed that the stent did not release significant left sirolimus after 28 days. There was consistency between the groups in the evaluations during the same time periods, which reflects that the stent has reliable standardization.

## Acknowledgement

The authors thank Invamed RD Global (Ankara, Turkey) for supporting this study.

## Contributions of authors

**HY** conceptualized the paper, analyzed, and interprated the data. **RD** searched the literature and wrote the article. **HY** supervised and edited. All authors have read and agreed to the final version of the manuscript.

## References

1. Roth GA, Mensah GA, Johnson CO, Addolorato G, Ammirati E, Baddour LM, et al. Global Burden of Cardiovascular Diseases and Risk Factors, 1990-2019: Update From the GBD 2019 Study. J Am Coll Cardiol. 2020;76(25):2982–3021. doi: 10.1016/j.jacc.2020.11.010.

2. Nusca A, Viscusi MM, Piccirillo F, De Filippis A, Nenna A, Spadaccio C, Nappi F, Chello C, Mangiacapra F, Grigioni F, Chello M, Ussia GP. In Stent Neo-Atherosclerosis: Pathophysiology, Clinical Implications, Prevention, and Therapeutic Approaches. Life (Basel). 2022 M8;12(3):393. doi: 10.3390/life12030393.

3. CDC. Coronary artery disease (CAD). Available at: https://www.cdc.gov/heartdisease/coronary_ad.htm. (Accesion date: April 19th, 2023)

4. Iqbal J, Serruys PW. Revascularization strategies for patients with stable coronary artery disease. J Intern Med. 2014;276(4):336–51. doi: 10.1111/joim.12243.

5. Xi T, Gao R, Xu B, Chen L, Luo T, Liu J, Wei Y, Zhong S. In vitro and in vivo changes to PLGA/sirolimus coating on drug eluting stents. Biomaterials. 2010;31(19):5151–8. doi: 10.1016/j.biomaterials.2010.02.003.

6. Coerkamp CF, Hoogewerf M, van Putte BP, Appelman Y, Doevendans PA. Revascularization strategies for patients with established chronic coronary syndrome. Eur J Clin Invest. 2022:e13787. doi: 10.1111/eci.13787.

7. Kawashima H, Zocca P, Buiten RA, Smits PC, Onuma Y, Wykrzykowska JJ, de Winter RJ, von Birgelen C, Serruys PW. The 2010s in clinical drug-eluting stent and bioresorbable scaffold research: a Dutch perspective. Neth Heart J. 2020;28(Suppl 1):78–87. doi: 10.1007/s12471-020-01442-w.

8. Buiten RA, Zocca P, von Birgelen C. Thin, very thin, or ultrathin-strut biodegradable or durable polymer-coated drug-eluting stents. Curr Opin Cardiol. 2020;35(6):705–711. doi:10.1097/HCO.0000000000000786.

9. Burzotta F, Brancati MF, Trani C, Porto I, Tommasino A, De Maria G, Niccoli G, Leone AM, Coluccia V, Schiavoni G, Crea F. INtimal hyPerplasia evAluated by oCT in de novo COROnary lesions treated by drug-eluting balloon and bare-metal stent (IN-PACT CORO): study protocol for a randomized controlled trial. Trials. 2012;13:55. doi: 10.1186/1745-6215-13-55.

10. Aliman O, Ardic AF. Usability, safety, efficacy, positioning of the Atlas drug-eluting coronary stent: Evaluation of preliminary perioperative results. Research Square. 2023; Preprint (Version 1). doi.org/10.21203/rs.3.rs-2728468/v1

11. Palmerini T, Biondi-Zoccai G, Stone GW. Stent selection to minimize the risk of stent thrombosis. Curr Opin Cardiol. 2014;29(6):578–85. doi: 10.1097/HCO.0000000000000102.

12. Schulz C, Herrmann RA, Beilharz C, Pasquantonio J, Alt E. Coronary stent symmetry and vascular injury determine experimental restenosis. Heart. 2000;83(4):462–467. doi:10.1136/heart.83.4.462

13. Bangalore S, Kumar S, Fusaro M, Amoroso N, Attubato MJ, Feit F, Bhatt DL, Slater J. Short- and long-term outcomes with drug-eluting and bare-metal coronary stents: a mixed-treatment comparison analysis of 117 762 patient-years of follow-up from randomized trials. Circulation. 2012;125(23):2873–91. doi: 10.1161/CIRCULATIONAHA.112.097014.

14. Xhepa E, Tada T, Cassese S, et al. Safety and efficacy of the Yukon Choice Flex sirolimus-eluting coronary stent in an all-comers population cohort. Indian Heart J. 2014;66(3):345–349. doi:10.1016/j.ihj.2014.05.003

15. Karahan O, Naci ÖC, Sümer T, Hafiz E, Khalil E. Investigation of the endothelial response of super elastic braided stent: An experimental evaluation. Acta Medica Alanya. 2020;4(3):236–41. doi.org/10.30565/medalanya.745576

16. Wilson GJ, Polovick JE, Huibregtse BA, Poff BC. Overlapping paclitaxel-eluting stents: long-term effects in a porcine coronary artery model. Cardiovasc Res. 2007;76(2):361–72. doi: 10.1016/j.cardiores.2007.07.004.

17. Norman G. Likert scales, levels of measurement and the “laws” of statistics. Adv Health Sci Educ Theory Pract. 2010;15(5):625–32. doi: 10.1007/s10459-010-9222-y.

18. Buszman P, Milewski K, Zurakowski A, et al. Novel biodegradable polymer-coated, paclitaxel-eluting stent inhibits neointimal formation in porcine coronary arteries. Kardiol Pol. 2010;68(5):503–509.

19. Shofti R, Zaretzki A, Cohen E, Engel A, Bar-El Y. The sheep as a model for coronary artery bypass surgery. Lab Anim. 2004;38(2):149–157. doi:10.1258/002367704322968821

20. Chan S. Targeting the mammalian target of rapamycin (mTOR): a new approach to treating cancer. Br J Cancer. 2004;91(8):1420–1424. doi:10.1038/sj.bjc.6602162

21. Chaabane C, Otsuka F, Virmani R, Bochaton-Piallat ML. Biological responses in stented arteries. Cardiovasc Res. 2013;99(2):353–363. doi:10.1093/cvr/cvt115

22. Joner M, Finn AV, Farb A, et al. Pathology of drug-eluting stents in humans: delayed healing and late thrombotic risk. J Am Coll Cardiol. 2006;48(1):193–202. doi:10.1016/j.jacc.2006.03.042

23. Buszman P, Trznadel S, Milewski K, et al. Novel paclitaxel-eluting, biodegradable polymer coated stent in the treatment of de novo coronary lesions: a prospective multicenter registry. Catheter Cardiovasc Interv. 2008;71(1):51–57. doi:10.1002/ccd.21392

24. Stone GW, Moses JW, Ellis SG, et al. Safety and efficacy of sirolimus- and paclitaxel-eluting coronary stents. N Engl J Med. 2007;356(10):998–1008. doi:10.1056/NEJMoa067193

25. Hansen KN, Jensen LO, Maeng M, et al. Five-Year Clinical Outcome of the Biodegradable Polymer Ultrathin Strut Sirolimus-Eluting Stent Compared to the Biodegradable Polymer Biolimus-Eluting Stent in Patients Treated With Percutaneous Coronary Intervention: From the SORT OUT VII Trial. Circ Cardiovasc Interv. 2023;16(1):e012332. doi:10.1161/CIRCINTERVENTIONS.122.012332

26. Rheude T, Koch T, Joner M, et al. Ten-year clinical outcomes of drug-eluting stents with different polymer coating strategies by degree of coronary calcification: a pooled analysis of the ISAR-TEST 4 and 5 randomised trials. EuroIntervention. 2023;18(14):1188–1196. doi:10.4244/EIJ-D-22-00781

27. Ijichi T, Nakazawa G, Torii S, et al. Late neointimal volume reduction is observed following biodegradable polymer-based drug eluting stent in porcine model. Int J Cardiol Heart Vasc. 2021;34:100792. doi:10.1016/j.ijcha.2021.100792

28. Park HK, Paick SH, Kim HG, Lho YS, Bae S. The impact of ureteral stent type on patient symptoms as determined by the ureteral stent symptom questionnaire: a prospective, randomized, controlled study. J Endourol. 2015;29(3):367–371. doi:10.1089/end.2014.0294

